# Previously undetected superspreading of *Mycobacterium tuberculosis* revealed by deep sequencing

**DOI:** 10.1101/801308

**Authors:** Robyn S. Lee, Jean-François Proulx, Fiona McIntosh, Marcel A. Behr, William P. Hanage

## Abstract

Tuberculosis disproportionately affects the Canadian Inuit. To address this, it is imperative we understand transmission dynamics in this population. We investigate whether ‘deep’ sequencing can provide additional resolution compared to standard sequencing, using a well-characterized outbreak from the Arctic (2011-2012, 50 cases). Samples were sequenced to ~500-1000x and reads were aligned to a novel local reference genome generated with PacBio SMRT sequencing. Consensus and heterogeneous variants were identified and compared across genomes. In contrast with previous genomic analyses using ~50x depth, deep sequencing allowed us to identify a novel super-spreader who likely transmitted to up to 17 other cases during the outbreak (35% of all cases that year). It is increasingly evident that within-host diversity should be incorporated into transmission analyses; deep sequencing can facilitate accurately detection of super-spreaders and corresponding transmission clusters. This has implications not only for TB, but all genomic studies of transmission - regardless of pathogen.

## Introduction

Tuberculosis (TB) in Canada is highest among the Inuit, an Indigenous population with a rate over 300x that of the non-Indigenous Canadian-born population in 2016. ^1^ Canada recently set a goal of TB elimination in the Inuit by 2030,^1^ which will not be achieved without halting ongoing transmission. Previous studies have used genomic data either alone or in conjunction with classical epidemiology to investigate TB transmission dynamics in the Canadian North,^2–4^ with the aim of identifying clusters to help guide public health interventions. Thus far, such studies have relied on identifying consensus single nucleotide polymorphisms (cSNPs), consistent with prevailing methodology in this field.

Recent studies suggest that incorporation of within-host diversity into genomic analyses may provide greater resolution of transmission than cSNP-based approaches alone.^5–8^ This may be particularly important for investigation of outbreaks occurring over short time scales and/or in settings such as the Canadian North, where the genetic diversity of circulating strains is especially low. In both of these circumstances, it is common to find many samples separated by zero cSNPs, hindering accurate source ascertainment. To investigate this hypothesis, we used deep sequencing (i.e., to ~10-fold more than standard, or 500-1000x) to re-evaluate transmission in a densely-sampled outbreak in Nunavik, Québec.

This outbreak, which has been previously described,^4,9^ comprised 50 microbiologically-confirmed cases of TB who were diagnosed in a single Inuit community between 2011-2012 - a rate of 5,359/100,000 for that year. Genomic epidemiology analyses using sequencing depths of ~50x that are standard in such work, identified multiple clusters of transmission in this outbreak, ^4^ however, there was insufficient genetic variation detected to infer precise person-to-person transmission events within these subgroups, given the short time frame and low mutation rate of *M. tuberculosis* (~0.5 SNPs/genome/year for Lineage 4 ^10^). In this study, we illustrate how within-host diversity can be incorporated into transmission analyses and in doing so, find new features of the transmission networks in this community, in particular, identifying a previously unrecognized superspreading event. We highlight a potential role for deep sequencing in public health investigations, with implications for TB control in Canada’s North as well as other high-transmission environments.

## Materials and methods

### Study subjects

All 50 samples from the 2011-2012 outbreak ^4^ were eligible for inclusion, as well as samples from all cases (n=15) diagnosed in same village in the preceding five years (2007 onwards), 13/15 of which were caused by the same strain of *M. tuberculosis* (the ‘Major [Mj]-III’ sublineage ^3^). There were two episodes of recurrent TB (i.e., where an individual had microbiologically-confirmed TB once, was cured, but developed TB again during the study period); otherwise, all samples are from unique individuals. All cases had pulmonary TB that was Lineage 4 (Euro-American ^4^). Cross-contamination was ruled out as described in ^4^.

### DNA extraction and sequencing

Laboratory methods are described in detail in the **Supplementary Material**. The Illumina HiSeq 4000 was used for paired-end 100bp sequencing. To obtain the target depth of coverage, pooled libraries were run on four independent lanes.

### Bioinformatics

Quality control of genomic data is described in detail in the **Supplemental Material**. Reads were aligned using the Burrows Wheeler Aligner MEM algorithm (v.0.7.15 ^11^) to the H37Rv reference (NC_000962.3 in the National Center for Biotechnology Information [NCBI] RefSeq database) and sorted using Samtools (v.1.5 ^12^). Analyses were later repeated using a local reference genome (described below). Reads with ambiguous mapping were excluded, as were reads with excessive soft-clipping (i.e., more than 20% of read length) based on our previous work. ^6^ Duplicate reads were marked using Picard MarkDuplicates (v.2.9.0, https://broadinstitute.github.io/picard/) and reads were locally re-aligned around indels using Genome Analysis ToolKit (GATK, v.3.8 ^13^). All sites were called using GATK’s Unified Genotyper algorithm.

Variants were filtered for quality using custom Python scripts (v.3.6) with the following thresholds: Phred < 50, Root Mean Squared Mapping Quality (RMS-MQ) ≤ 30, depth (DP) < 20, Fisher Strand Bias (FS) ≥ 60 and read position strand bias (ReadPos) < −8. ^6^ cSNPs were classified as positions where ≥ 95% of reads were the alternative allele (ALT), hSNPs were classified as positions where > 5% and < 95% of reads were ALT, and positions with the ALT present in < 5% of reads were classified as ‘reference’. We also compared inferences of transmission from this analysis to i) when these thresholds were increased to the minimum values among cSNPs in the initial H37Rv analysis, and ii) when cSNPs were classified using a threshold of ≥ 99%, and hSNPs were classified when 1% < ALT < 99%, in order to assess the robustness of inferences to different filtering protocols.

Low-quality variants, variants in proline-proline-glutamic acid (PE) and proline-glutamic-acid/polymorphic-guanine-cytosine-rich sequence (PE_PGRS) genes, transposons, phage and integrase, and positions with missing data, were excluded. All samples were drug-susceptible, except for MT-6429, which was rendered resistant to isoniazid by a frameshift deletion at position 1284 in the catalase-peroxidase gene *katG*. As such, positions associated with drug resistance were not masked in this analysis.

Concatenated cSNP alignments were generated excluding positions with hSNPs. Pairwise cSNP distances between samples were computed using snp-dists (v.0.6, available at https://github.com/tseemann/snp-dists). The frequency of hSNPs at each position in the genome was tabulated and hSNPs were reviewed to identify variants shared between samples.

### Phylogenetics and clustering

Core cSNP alignments were used to generate maximum likelihood trees using IQ-Tree (v.1.6.8 ^14^). Model selection was based on the lowest Bayesian Information Criterion. Hierarchical Bayesian Analysis of Population Structure ^15^ was run in R (v.3.5.2) to identify clusters. See the **Supplementary Material** for additional detail.

### Single Molecule Real-Time (SMRT) sequencing and assembly

To examine the influence of potential alignment errors in identification of hSNPs, we used SMRT sequencing with the PacBio RSII platform to create a local reference genome for the outbreak. Sample MT-0080 was chosen for sequencing because this was previously identified as the probable source for as many as 19 of the 50 cases diagnosed in 2011-2012.^4^ A single colony from the culture was selected for SMRT sequencing and Illumina MiSeq (for polishing of the long-read assembly). Further detail is provided in the **Supplementary Material**.

Long-reads were assembled and corrected using Canu (v.1.7.1 ^16^). Pilon (v.1.23 ^17^) was then used to polish the assembly and was re-run until no further corrections were possible. Quast (v.5.0.2, ^18^) was used to evaluate assembly quality. RASTtk (v.2.0 ^19^) was used for annotation, to identify regions for masking as previous.

### Epidemiological data

Epidemiological and clinical data were collected on all cases and contacts using standardized questionnaires, as part of the routine public health response.

### Statistical analyses

A two-sample test of proportions was used to compare overall proportions across references, and the Wilcoxon Signed Rank test was used to compare paired SNP distances. Analyses were done in Stata (v.15, StataCorp, College Station, TX, USA).

### Data availability

Sequencing data and the assembly for MT-0080 are available on the NCBI’s Sequence Read Archive under BioProject PRJNA549270.

### Ethics

Ethics approval was obtained from the Institutional Review Board (IRB) of the Harvard T.H. Chan School of Public Health (IRB18-0552) and the IRB of McGill University Faculty of Medicine (IRB A02-M08-18A). All data was analyzed in non-nominal fashion. This study was done with approval of and in collaboration with the Nunavik Regional Board of Health and Social Services.

## Results

62/65 (95·4%) available TB samples from cases diagnosed between 2007-2012 were successfully sequenced and passed quality control. This included 48/49 (98·0%) of the samples with an identical Mycobacterial Interspersed Repetitive Units Variable Number Tandem Repeats (MIRU-VNTR) pattern during the outbreak year. The remaining three samples could not be re-grown. Reads that were non-MTBC were removed (**Table S1**) and there was no obvious association between percent contamination and hSNP frequency. Epidemiological and clinical data on all outbreak cases are described in ^4^.

Average genome coverage and depth across the H37Rv reference was 98·64% [SD 0·07%] and 714·53 [SD 92·68], respectively. Our primary filtering protocol yielded 51,430 cSNPs and 4,897 hSNPs across all individual samples (**Table S2**). Excluding positions that were invariant compared to the reference or where any sample was missing and/or was low-quality resulted in a core alignment of 860 cSNP positions and 136 hSNP positions (note, these are not mutually exclusive, as positions with cSNPs in some samples may have hSNPs in others).

42 positions had hSNPs that were shared across all 62 samples (**Table 1**, **Supplementary Dataset 1A**). Depth of coverage at these positions was, on average, 39% higher than the average depth across the same sample (SD 36·7%, **Supplementary Dataset 1B**). Along with manual review of alignments (**Figure S1)**, this suggested that many of these were false positives, potentially due to alignment error (e.g., from underlying structural variation in our samples compared to the H37Rv reference).

**Table 1.** Comparison of alignments to H37Rv and MT-0080_PB

|  | H37Rv (4,411,532 bp) | MT-0080_PB (4,426,525 bp) | P value |
| --- | --- | --- | --- |
| **Number of positions according to reference genome** |  |  |  |
| Invariant reference across all samples, n (%) | 4,018,786 (91.10%) | 4,084,195 (92.27%) | < 0.00005 <sup>a</sup> |
| Position was missing / low quality in at least one sample, n (%) | 391,761 (8.88%) | 342,179 (7.73%) | < 0.00005 <sup>a</sup> |
| Position was an c/hSNP in at least one sample, n (%) | 985 (0.22%) | 152 (0.00%) | < 0.00005 <sup>a</sup> |
| Shared cSNPs across all samples, n (%) | 764 (0.02%) | 1 (0.00%) | < 0.00005 <sup>a</sup> |
| Shared hSNPs across all samples, n (%) | 42 (0.00%) | 0 (0%) | < 0.00005 <sup>a</sup> |
| Core pairwise distances |  |  |  |
| Core cSNPs vs. reference, median (range) | 791 (790-792) | 3 (1-65) | < 0.00005 <sup>b</sup> |
| Core cSNPs between samples, median (range) | 3 (0-64) | 3 (0-66) | < 0.00005 <sup>b</sup> |

To address this, we generated a local reference genome for the outbreak, MT-0080_PB. Quality metrics for the MT-0080_PB assembly are given in **Table S3**. Compared to H37Rv, mean genome coverage and depth were higher with MT-0080_PB (at 99·33% [SD 0·09%] and 717·07 [SD 93·01], respectively), fewer positions were missing/low-quality (p < 0·00005, **Table 1**), and overall, fewer variable positions were detected (p < 0·00005). While core cSNP distances were similar between samples regardless of the reference (**Table 1**), the number of hSNPs was greatly reduced using MT-0080_PB (**Table S2**); while 4,897 hSNPs were identified across all individual samples using H37Rv, only 125 hSNPs were identified using MT-0080_PB. There were also no hSNPs shared across all 62 samples using MT-0080_PB. Together, these findings support our hypothesis that alignment error is responsible for many of the detected variants, and indicate a local reference is important for accurate identification of hSNPs. All further results presented are based on the MT-0080_PB alignment.

A maximum likelihood tree was generated from 94 core cSNP positions (excluding sites invariant across all samples and the reference) compared to MT-0080 (**Figure 1A)**. Consistent with previous work,^4^ hierBAPS identified two main sub-lineages (‘Mj-V’ and ‘Mj-III’ per ^3^), with three sub-clusters (Mj-IIIA/B/C).

**Figure 1.**
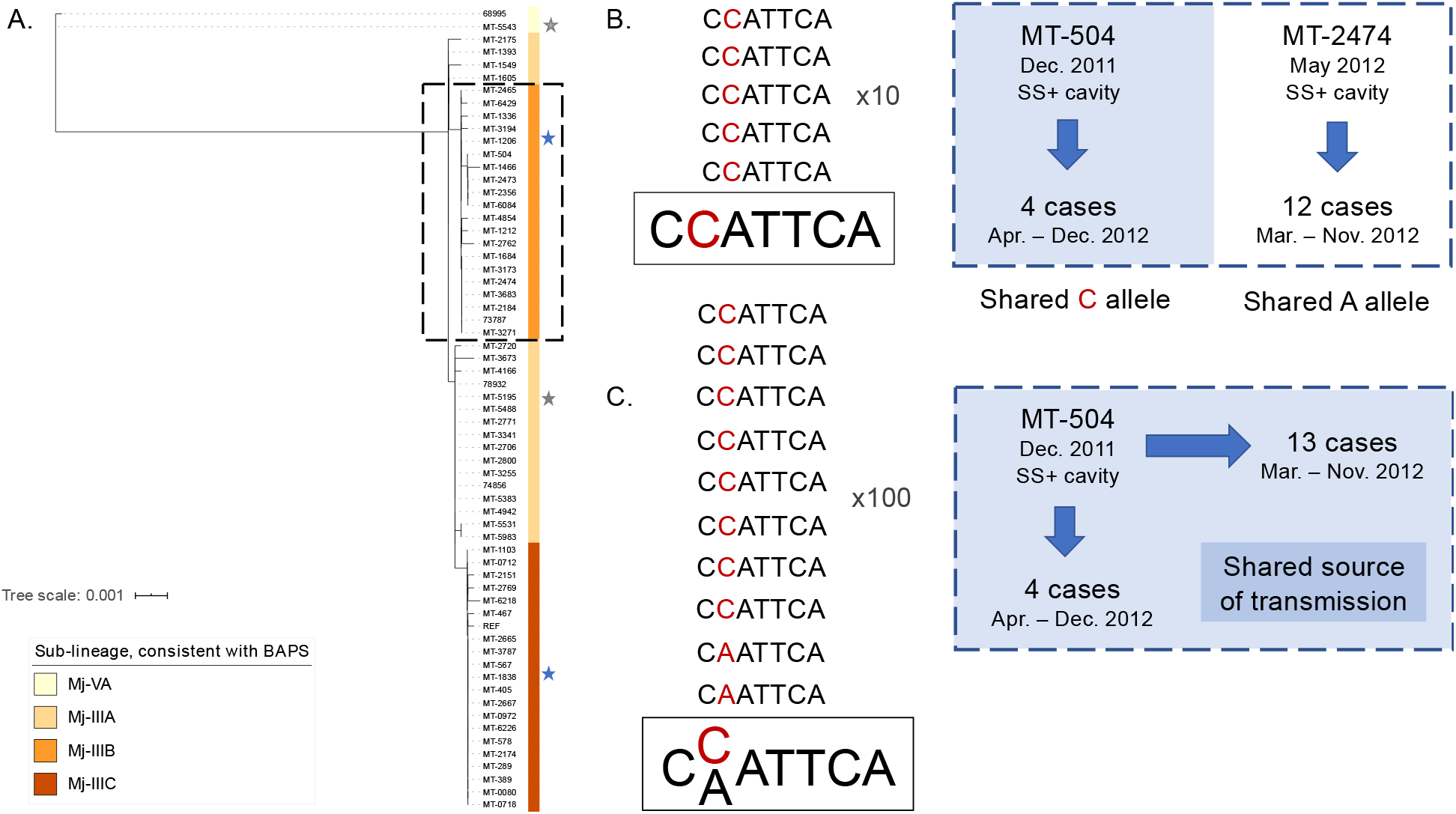
Transmission of *M. tuberculosis* in village K. *Panel A.* Maximum likelihood tree of 62/65 cases diagnosed between 2007-2012 in village K based on consensus single nucleotide polymorphisms (cSNPs). After aligning to a local reference, MT-0080_PB, cSNPs were identified based on a minimum threshold of ≥ 95% of reads supporting the alternative allele. A core cSNP alignment was then produced with 103 positions. IQ-Tree (v.1.6.8 ^14^) was then used to generate the tree using a KP3 model with correction for ascertainment bias. Model selection was based on the lowest Bayesian Information Criterion. Clusters were identified using hierarchical Bayesian Analysis of Population Structure.^15^ These clusters were consistent with the sub-lineages previously identified in ^3,4^, thus only sub-lineage names are indicated. During this time period, there were two individuals who had a second episode of TB; stars are used to highlight these samples, with a different colour for each patient. MT-0080 is included in the alignment as the deep sequencing data from a sweep of all colonies identified a cSNP compared to the MT-0080_PB reference, which itself was generated from a single colony pick. *Panel B*. Standard sequencing (to ~40-50x), along with epidemiological data, had indicated that the Major [Mj]-IIIB sub-lineage was comprised of two subgroups of five and 13 patients, respectively.^4^ MT-504 was the suspected source case for the subgroup of five, which all shared a ‘C’ allele at position 276,685 in H37Rv (position 276,544 in MT-0080_PB). In contrast, all members of the subgroup of 13 shared an ‘A’ at this position. Previously, MT-2474 was the suspected source case for this subgroup; this case was the first person with smear-positive (SS+) cavitary disease diagnosed in this subgroup. Panel C. In contrast to standard sequencing, deep sequencing data revealed that, in fact, MT-504 – the presumed source for the subgroup of five cases and the first highly contagious case diagnosed in Mj-IIIB during the outbreak year – had both ‘C’ and ‘A’ alleles at this position (563:133 of reads, respectively), suggesting this was in fact the most probable source for both subgroups.

### hSNPs identify super-spreaders and more accurately resolve transmission clusters

The core cSNPs and hSNPs between samples are shown in **Supplemental Dataset 2A**, with the sub-groups identified in the original analysis indicated. Overlaying hSNPs with the cSNP-based analysis revealed a novel super-spreader (MT-504) in Cluster Mj-IIIB, undetected by genomic epidemiology analyses relying on lower sequencing depth. ^4^ MT-504 had smear-positive cavitary disease and was diagnosed in late 2011; previous analyses had found this case shared a single cSNP with four other cases diagnosed from March – December of 2012 (position 276,685 according to H37Rv / 276,544 in the MT-0080_PB alignment, **Supplementary Dataset 2**). Coupled with epidemiological data on contact (shared attendance at local community ‘gathering houses’, social venues specifically identified by public health during the outbreak), this strongly supported transmission from MT-504 to other members of this subgroup. In contrast, the other subgroup of Mj-IIIB with 13 cases did not share this cSNP. This initially refuted transmission, as we would expect 0 SNPs to accrue in recent transmission given the short time period, low mutation rate of TB, and overall low diversity of strains circulating in the village (**Figure 1B**). Instead, we previously postulated that the first smear-positive case in this subgroup (MT-2474, diagnosed in May 2012) led to the majority of transmission (note, the first smear-negative case in this subgroup was diagnosed in March 2012). However, deep sequencing data suggest otherwise; these data show that MT-504 harboured both the reference (133 reads [19·1%]) allele, present in the subgroup of 13, as well as the alternative allele (563 reads [80·9%]) at this position (**Figure 1C**). As MT-504 was the first contagious case diagnosed in Mj-IIIB, and all 13 cases in this subgroup had attended or resided in a gathering house (with 9/13 [69·2%] reporting attendance at the same houses as MT-504), this strongly suggests that MT-504 is in fact the most probable source for both subgroups.

### hSNP analysis adds support for suspected transmission

Sample 68995 and MT-5543 were from 2007, and were the only strains from the Mj-VA sub-lineage in this village. Previous analysis indicated Mj-VA strains from other villages were distantly related,^4^ while these two samples were separated from one another by zero core cSNPs. This suggests direct transmission between these historical cases, a hypothesis strongly supported by hSNP analysis, as the samples share hSNPs that are not found in any other sample in the dataset. These hSNPs were present even when highly conservative filtering thresholds were used **(Supplemental Dataset 2B**). Importantly, these hSNPs were not detected when using H37Rv as the reference.

### Potential utility for discriminating TB recurrence

Six individuals had TB recurrence in 2011-2012. Paired samples were available for two of these (Patient 1: samples MT-5195 in 2007 and MT-1838 in 2012; Patient 2: samples MT-5543 in 2007 and MT-1206 in 2012, **Figure 1A**). cSNP-based analyses suggested their second episodes of TB were due to re-infection with a new strain, rather than relapse with the strain causing their original disease. Investigation of within-host diversity strongly supported this conclusion; using deep sequencing, we verified that there was a single, different strain present at both baseline and their second episodes of TB. There was no evidence for mixed infection at either baseline or second episode with these strains, more definitively ruling out relapse in this low diversity setting (**Supplemental Dataset 2A/B/C**).

### Impact of altering cSNP and hSNP thresholds

To ensure we were not missing lower frequency variants using the prior cSNP/hSNP thresholds, we re-ran our analysis such that hSNPs were classified when 1% < ALT < 99%. Quality scores for individual cSNPs and hSNPs are given in **Table S4** and the core cSNP/hSNP alignment is shown in **Supplemental Dataset 2C**. While our primary analysis using a threshold of > 95% for cSNPs identified a single cSNP (A>G) shared across all samples compared to MT-0080_PB, close examination of the MT-0080 deep sequencing data (obtained using DNA from a sweep of the plate) showed that this sample had both alleles at this position, with only the minority ‘A’ allele (33 reads/1189 [2·8%]) isolated for SMRT sequencing. Based on this, we recommend sequencing samples both using a clean sweep (with an alternative sequencing platform) and a single colony pick when generating a reference genome for TB, as using the latter alone may introduce error and affect epidemiological inferences. With this exception, no other informative hSNPs were detected using these thresholds.

## Discussion

As the TB epidemic continues among the Canadian Inuit, targeted public health interventions are essential to halt ongoing transmission. In order to do so, it is important that transmission events and associated risk factors are accurately identified. Our previous work suggested that hSNP analysis could enhance resolution of TB transmission ^6^. To investigate how this approach could be applied for TB control, we used deep sequencing to re-examine a major TB outbreak in the Canadian Arctic.

Several recent studies, including work by the authors^6^, have shown that *M. tuberculosis* within-host diversity can be transmitted between individuals^8,20^. Using deep sequencing data allowed us to better identify this diversity in a Nunavik outbreak compared to previous analyses with standard sequencing depth, ^3,4^ and facilitated detection of a novel super-spreader who was likely responsible for ~1/3 of the cases from 2011-2012. Super-spreading has been described in a number of pathogens,^21^ including TB.^22^ Our findings suggest this can play an important role in driving TB outbreaks. We therefore propose that investigation of within-host diversity is necessary to ensure detection of such super-spreaders in this context, and potentially other high-transmission environments as well, in order to accurately identify transmission networks and associated risk factors. We typically only know that a person is a super-spreader retrospectively, i.e., once they have already transmitted to many others and once corresponding genomic data is available. However, if we can identify the characteristics associated with this phenomenon at the population level, this could be used prospectively to predict whether cases are likely to be super-spreaders - as they are diagnosed. This could allow resources to be better allocated, for example, investigation of their contacts could be prioritized and/or targeted screening of the social venues they attend could be rapidly initiated, potentially leading to faster detection of secondary cases and initiation of prophylaxis for new infections. In the case of MT-504, nearly all of the secondary cases had attended the same local community gathering houses as the putative source; this also strongly suggests the importance of these venues in facilitating transmission in this setting.

Several studies have used genomics to investigate TB recurrence,^23–25^ however, the methods used to assess for mixed infection at either time point have been inconsistent and may not be sufficient to discriminate recurrence in settings with low strain diversity. In this analysis, we provide proof-of-principle that deep sequencing can potentially help rule out relapse. The distinction between relapse and re-infection is important at individual and population levels; high rates of relapse in a community would indicate a problem with treatment or adherence, potentially warranting changes to clinical management, while re-infection would indicate the need for public health interventions such as activate case finding. Also, individuals in Nunavik who have had prior treatment for active TB disease in the past are also not routinely offered prophylaxis on re-exposure, based on historical data suggesting ~80% protection is afforded by prior infection.^26^ The degree to which re-infection drives recurrence in Nunavik is currently unknown, but if re-infection is the primary cause, this clinical practice may need to be re-evaluated. Population-level genomic studies are currently underway to evaluate this.

To use deep sequencing to investigate within-host diversity, it is critical we minimize false positive hSNPs. We have shown that using a local strain as a reference not only reduces error, but improves detection of epidemiologically-informative variants. Genomic differences between outbreak strains and H37Rv have been previously illustrated by ^27,28^, with O’Toole *et al.* ^28^ warning that clinical TB strains may be needed to fully detect virulence genes in reference-based analyses. We propose these are also warranted for hSNP analysis. Where possible, we recommend using long-read sequencing to generate complete and local reference genomes.

Overall, our study has a number of strengths. Firstly, we had access to a densely-sampled outbreak, which was previously investigated using ‘standard’ sequencing depth and for which detailed epidemiological data was available. This allowed us to readily compare methodological approaches, showing how and when deep sequencing might be beneficial for public health. In doing this, we have identified important methodological considerations for hSNP detection, with implications for transmission analyses, but also potentially for resistance prediction as well.^29^ Finally, the use of long-read data has allowed us to completely assemble a novel TB genome from Nunavik. This will serve as a valuable resource for future studies of transmission in Nunavik (given the low strain diversity in the region ^3^), as well as other Inuit territories.

A potential limitation of this work is that, given the historical nature of the outbreak, deep sequencing was done using DNA extracted from culture. Due to methodological challenges of sequencing directly from sputum,^30–32^ few studies have examined the effect of culture on genome diversity. A recent study by Shockey *et al*^33^. using cSNPs suggests that variation may be lost during the culturing process, however authors did not examine the impact on hSNPs or transmission, and several other studies ^31,32,34^ previously found congruent results between cSNP analyses from culture versus raw samples. In terms of hSNPs, Votintseva *et al.* found no difference in the number detected between approaches.^31^ While ^32,34^ reported detecting fewer hSNPs with sequencing from culture versus from sputum, in ^34^, the median hSNPs was only 4.5 versus 5 hSNPs, respectively – a difference that may not be clinically significant, regardless of statistical significance. Given the inconsistency of results and paucity of data, further study is needed to understand how hSNP diversity may be affected by the culturing process, and to assess whether this affects inferences of transmission. We note that it is likely that enhanced detection of the hSNPs present in sputum would improve the resolution over that which we present in this work.

Another potential limitation is that, while we can compare the epidemiological inferences made between our previous analysis and our deep sequencing analysis, the bioinformatics pipelines themselves are not directly comparable. Methods to accurately identify hSNPs and incorporate them into transmission analyses are currently an area of active research. We illustrated in our recent paper ^6^ that additional steps and strict thresholds must be used to minimize false positive hSNPs, and have conducted the current analysis in consideration of this. However, we note that pre-filtering, our 2015 analysis found that MT-504 had five reference alleles at position 276,685 in the H37Rv alignment (out of 75) and randomly downsampling the current data to simulate ~50x yielded similar results (5/47 reads at position 276,544 aligned to MT-0080_PB). As most genomic studies of TB employ conservative thresholds of 75-90% allele frequency to classify cSNPs, many bioinformatics pipelines would consider this heterogeneity as potentially suspect at standard sequencing depth. This therefore reinforces the need for greater depth and/or analytic approaches (e.g., ^35^) to ensure accurate discrimination of sequencing/analytic error from true variation.

In summary, we have found evidence of mixed variants with important epidemiologic implications that would not have been detected with standard methods and common filtering criteria. This illustrates that genomic methods, while powerful, still require careful interpretation and can still harbor considerable ambiguity when comparing very close links in a transmission chain – a finding whose relevance likely extends far beyond TB, given the increasing number of pathogens undergoing genomic investigation. We demonstrate that deep sequencing can aid in transmission analyses, in particular by allowing accurate identification of TB super-spreaders and key associated epidemiological characteristics. In terms of TB control, this work has important implications for the Canadian North as well as other regions with high TB transmission; as next-generation sequencing becomes a mainstay in public health surveillance, it is critical we recognize the limitations of analyses done using routine sequencing data and consider where and when deep sequencing might be warranted. Although costs continue to decline, we recognize deep sequencing of all samples in an outbreak may not be economically viable for every public health unit. As such, we propose that public health units using routine sequencing for tuberculosis consider – at a minimum – targeted deep sequencing of the more contagious (e.g., smear-positive) cases, in lieu of all samples, to help ensure accurate identification of super-spreaders and clusters of transmission. This may help TB control programmes better understand the risk factors for such transmission and enable prioritization of public health resources in future outbreaks.

## Acknowledgements

We would like to thank the council of the village for their ongoing support of this work. We also acknowledge staff from the Centre Locales des Services Communautaires and the Nunavik Regional Board of Health and Social Services for their hard work during the outbreak. We would like to thank Dr. Hafid Soualhine (currently at the National Reference Centre for Mycobacteriology in the National Microbiology Laboratory, Public Health Agency of Canada) for previous confirmation of the frameshift deletion in *katG* of MT-6429. We also thank Dr. Anders Gonçalves da Silva and Dr. Glen Carter of the Microbiological Diagnostic Unit Public Health Laboratory at the University of Melbourne for their helpful input on high-quality SMRT sequencing. Library preparation and sequencing for all samples was done at the Genome Québec / McGill Innovation Centre and high-performance computing was done using the Odyssey cluster from the Faculty of Arts and Science, Harvard University. This work was supported by an R01 grant from the National Institutes of Health, awarded to WPH (R01AI128344). RSL is also supported by a Fellowship from the Canadian Institutes of Health Research (MFE 152448). MAB holds a Canadian Institutes of Health Research Foundation Grant (FDN-148362).

## Contributions

RSL and WPH conceived and designed the study. RSL designed and did the analyses, made the tables and figures, interpreted the data, and wrote the first draft of manuscript. JFP provided epidemiological data. MAB provided the bacterial samples, as well as laboratory reagents, Biosafety Level 3 access and technician labour in-kind to RSL. FM did the culture and DNA extraction for the HiSeq and PacBio SMRT sequencing. WPH reviewed the initial draft. All authors provided feedback and reviewed the final version of the manuscript before submission. RSL had full access to the data and final responsibility for the decision to submit for publication.

