## Supplementary_Material for "Previously undetected superspreading of *Mycobacterium tuberculosis* revealed by deep sequencing"

**deep sequencing**

Robyn S. Lee PhD^1,2^*, Jean-François Proulx MD^3^, Fiona McIntosh BSc^4^, Marcel A. Behr MD^4^,

William P. Hanage PhD^1,2^

1. Center for Communicable Disease Dynamics, Harvard T. H. Chan School of Public Health, 677 Huntington Avenue, Kresge Building, Room 506I, Boston, MA, USA, 02115
2. Department of Epidemiology, Harvard T. H. Chan School of Public Health, Boston, MA, USA, 677 Huntington Avenue, Kresge Building, 5^th^ floor, Boston, MA, USA, 02115
3. Nunavik Regional Board of Health and Social Services, Kuujjuaq, Québec, Canada
4. The Research Institute of McGill University Health Centre, Montréal, Québec, Canada

**Supplementary Material**

**Contents**

- Supplementary Methods
- Supplementary Figure S1
- Supplementary Tables S1-S4
- Supplementary Datasets 1 and 2

**Supplementary Methods.**

***Study subjects.*** All samples from the 2011-2012 ‘outbreak’ (50 microbiologically-confirmed cases, one sample per case) were included, as well as samples from all cases (n=15) diagnosed in same village in the five years preceding the outbreak (2007 onwards), 13/15 of which were caused by the same strain of *M. tuberculosis* (the ‘Major [Mj]-III’ sublineage ^1^). All were Lineage 4 (Euro-American ^2^) and all had pulmonary TB. Cross-contamination between samples was previously ruled out, as described.^2^

***DNA extraction and sequencing with Illumina HiSeq.*** Samples were cultured once on Middlebrook 7H10 agar and plate sweeps were collected for DNA extraction using the van Soolingen method.^3^ Genomic DNA was quantified using the Quant-iT PicoGreen dsDNA Assay (ThermoFisher Scientific, Massachusetts, USA). Library preparation and sequencing were done at the McGill University/Genome Québec Innovation Centre. The Illumina HiSeq 4000 was used to produce paired-end 100bp reads. To obtain the depth of coverage needed for this study (~500-1000x for deep sequencing, compared to ~50-100x as routinely done by public health), pooled libraries were run on four independent lanes.

***Quality control of genomic data.*** FastQC (v.0.11.5, https://www.bioinformatics.babraham.ac.uk/projects/fastqc/) was used to assess sequencing data quality and reads were trimmed to remove low-quality bases using Trimmomatic (v.0.36 ^4^). Kraken (v.1.1 ^5^) was then used to identify potential contamination with the miniKraken database (minikraken_20171019). Reads classified as ‘Mycobacterium tuberculosis complex’ were extracted using Seqtk (v.1.2, Li H, available at: https://github.com/lh3/seqtk).

***Reference-based assembly and identification of variants.*** Reads were first aligned using the Burrows Wheeler Aligner MEM algorithm (v.0.7.15 ^6^) to the H37Rv reference (NC_000962.3 in the National Center for Biotechnology Information [NCBI] RefSeq database) and sorted using Samtools (v.1.5 ^7^). Analyses were later repeated using a local reference genome (described below). Reads were with ambiguous mappings were excluded, as were reads with excessive soft-clipping (i.e., more than 20% of read length) based on our previous work.^8^ Duplicate reads were marked using Picard MarkDuplicates (v.2.9.0, https://broadinstitute.github.io/picard/) and reads were locally re-aligned around indels using Genome Analysis ToolKit (GATK, v.3.8 ^9^). All sites were called using GATK’s Unified Genotyper algorithm under a diploid model, with the -d 1500 to avoid down-sampling to 250 (done by default with this tool during variant calling). Variants (cSNPs and heterogenous SNPs [hSNPs]) versus H37Rv were annotated using snpEff (v.4.3t ^10^).

Variants were filtered for quality using custom Python scripts (v.3.6) with the following thresholds: Phred < 50, Root Mean Squared Mapping Quality (RMS-MQ) ≤ 30, depth (DP) < 20, Fisher Strand Bias (FS) ≥ 60 and read position strand bias (ReadPos) < -8.^8,11^ cSNPs were classified as positions where ≥ 95% of reads were the alternative allele (ALT), hSNPs were classified as positions where > 5% and < 95% of reads were ALT, and positions with the ALT present in <5% of reads were classified as ‘reference’. We also compared inferences of transmission from this analysis to i) when thresholds were increased to the minimum values among cSNPs in the initial H37Rv analysis, and ii) when cSNPs were classified using a threshold of ≥ 99%, and hSNPs were classified when 1% < ALT < 99%, in order to assess the robustness of inferences to different filtering protocols.

Full genome alignments were generated for each analysis. Low-quality variants, variants in proline-proline-glutamic acid (PE) and proline-glutamic-acid/polymorphic-guanine-cytosine-rich sequence (PE_PGRS) genes, transposons, phage and integrase, and positions with missing data, were excluded. All samples were drug-susceptible, except for MT-6429, which was resistant to isoniazid due to a frameshift deletion at position 1284 in *katG*. As such, positions associated with drug resistance were not masked in this analysis. Alignments with informative hSNPs were reviewed using Tablet (v.1.17.08.17, ^15^).

Concatenated cSNP alignments using snp-sites -c (v.2.4.0 ^12^), with positions with hSNPs excluded. Pairwise cSNP distances between samples were computed using snp-dists (v.0.6, available at https://github.com/tseemann/snp-dists). The frequency of hSNPs at each position in the genome was tabulated and hSNPs were reviewed to identify variants shared between samples.

***Phylogenetics and clustering.*** Concatenated core cSNP alignments were used to generate maximum likelihood trees using IQ-Tree (v.1.6.8 ^13^). Model selection was based on the lowest Bayesian Information Criterion. Hierarchical Bayesian Analysis of Population Structure ^14^ was run in R (v.3.5.2) using RhierBAPS ^15^ to identify clusters. Phylogenetic trees were visualized using Interactive Tree of Life.^16^

***Single Molecule Real-Time (SMRT) sequencing and assembly.*** To examine the influence of potential alignment errors in identification of hSNPs, we used SMRT sequencing with the PacBio RSII platform to create a local reference genome. Sample MT-0080 was chosen for sequencing because this was previously identified as the probable source for as many as 19 of the 50 cases diagnosed in 2011-2012.^2^ Prior to sequencing, the culture was grown on a Middlebrook 7H10 agar plate. A single colony was then selected and grown further in 3mL of Middlebrook 7H9 Broth to provide sufficient DNA for SMRT sequencing and Illumina MiSeq (for polishing of the long-read assembly). DNA for SMRT sequencing was extracted using the MagAttract High Molecular Weight DNA Kit from Qiagen (Maryland, USA). High molecular weight fragments were verified using gel electrophoresis. Library preparation and sequencing were then done at the McGill University/Genome Québec Innovation Centre. Prior to sequencing, fragment size was evaluated using a BioAnalyzer and the BluePippin system (Sage Science, Massachusetts, USA) was used for size selection. DNA for Illumina MiSeq was extracted using the van Soolingen method, as previous.^3^

Long-reads were assembled and corrected using Canu (v.1.7.1 ^17^). Pilon (v.1.23 ^18^) was then used to polish the assembly and was re-run until no further corrections were possible. Quast (v.5.0.2 ^19^) was used to evaluate assembly quality and RASTtk (v.2.0 ^20^) was used for annotation. Average nucleotide identity (in percent) compared to the H37Rv reference was calculated using jSpecies (v.1.2.1 ^21^) with the MUMmer algorithm (v.3.23 ^22^), following the approach used in ^11,23^.

***Epidemiological data.*** Detailed epidemiological and clinical data was collected by the local healthcare team and the regional public health unit, as described in ^2,24^.

***Statistical analyses.*** A two-sample test of proportions was used to compare overall proportions across references, and the Wilcoxon Signed Rank test was used to compare paired SNP distances. Analyses were done in Stata (v.15, StataCorp, College Station, TX, USA).

***Data availability.*** Illumina HiSeq and PacBio SMRT sequencing data are available on the NCBI’s Sequence Read Archive under BioProject PRJNA549270, along with the complete assembly for MT-0080.

***Ethics****.* Ethics approval was obtained from the Institutional Review Board (IRB) of the Harvard T.H. Chan School of Public Health (IRB18-0552) and the IRB of McGill University Faculty of Medicine (IRB A02-M08-18A). All data was analyzed in non-nominal fashion. This study was done with approval of and in collaboration with the Nunavik Regional Board of Health and Social Services.

**Tables**

**Table S1.** Percent of kmers classified as *Mycobacterium tuberculosis* Complex with hSNP frequency after removing these, reported for the alignment to MT-0080_PB and these filtering thresholds: Phred score < 50, Root Mean Square Mapping Quality [RMS-MQ] ≤ 30, depth [DP] < 20, Read Position Rank Sum [ReadPosRankSum] < -8, Fisher Strand Bias [FS] ≥ 60

| Sample | Median percent MTBC across lanes | Minimum percent MTBC across lanes | Maximum percent MTBC across lanes | Frequency of hSNPs identified after removing non-MTBC |
| --- | --- | --- | --- | --- |
| 68995 | 98.81 | 98.67 | 98.85 | 1 |
| 73787 | 98.77 | 98.63 | 98.81 | 1 |
| 74856 | 98.75 | 98.58 | 98.8 | 0 |
| 78932 | 98.75 | 98.61 | 98.78 | 3 |
| MT-0080 | 98.80 | 98.65 | 98.84 | 1 |
| MT-0712 | 98.74 | 98.56 | 98.78 | 0 |
| MT-0718 | 98.53 | 98.37 | 98.57 | 5 |
| MT-0972 | 96.99 | 96.84 | 97.04 | 0 |
| MT-1103 | 98.57 | 98.41 | 98.62 | 2 |
| MT-1206 | 98.62 | 98.49 | 98.67 | 4 |
| MT-1212 | 98.55 | 98.41 | 98.59 | 4 |
| MT-1336 | 98.47 | 98.3 | 98.52 | 6 |
| MT-1393 | 98.66 | 98.53 | 98.7 | 2 |
| MT-1466 | 98.60 | 98.42 | 98.64 | 0 |
| MT-1549 | 98.58 | 98.44 | 98.62 | 1 |
| MT-1605 | 98.60 | 98.48 | 98.65 | 3 |
| MT-1684 | 98.47 | 98.33 | 98.52 | 0 |
| MT-1838 | 98.77 | 98.61 | 98.8 | 0 |
| MT-2151 | 98.65 | 98.53 | 98.69 | 0 |
| MT-2174 | 98.57 | 98.43 | 98.62 | 3 |
| MT-2175 | 98.56 | 98.43 | 98.6 | 2 |
| MT-2184 | 98.47 | 98.33 | 98.51 | 4 |
| MT-2356 | 98.40 | 98.22 | 98.44 | 5 |
| MT-2465 | 98.15 | 98.02 | 98.2 | 3 |
| MT-2473 | 98.56 | 98.44 | 98.6 | 2 |
| MT-2474 | 98.57 | 98.42 | 98.61 | 0 |
| MT-2665 | 97.68 | 97.28 | 97.74 | 2 |
| MT-2667 | 98.45 | 98.29 | 98.5 | 7 |
| MT-2706 ^a^ | 98.77 | 98.56 | 98.78 | 1 |
| MT-2720 | 98.53 | 98.4 | 98.58 | 11 |
| MT-2762 | 98.56 | 98.41 | 98.6 | 4 |
| MT-2769 | 98.45 | 98.32 | 98.49 | 4 |
| MT-2771 | 98.74 | 98.56 | 98.79 | 0 |
| MT-2800 | 98.70 | 98.54 | 98.75 | 2 |
| MT-289 | 98.70 | 98.52 | 98.74 | 1 |
| MT-3173 | 97.92 | 97.54 | 97.97 | 0 |
| MT-3194 | 98.60 | 98.46 | 98.64 | 2 |
| MT-3255 | 98.74 | 98.57 | 98.79 | 0 |
| MT-3271 | 98.51 | 98.34 | 98.56 | 1 |
| MT-3341 | 98.80 | 98.66 | 98.85 | 3 |
| MT-3673 | 98.82 | 98.69 | 98.85 | 0 |
| MT-3683 | 98.76 | 98.59 | 98.79 | 0 |
| MT-3787 | 98.83 | 98.69 | 98.86 | 0 |
| MT-389 | 98.59 | 98.44 | 98.63 | 1 |
| MT-405 | 98.75 | 98.57 | 98.78 | 0 |
| MT-4166 | 98.50 | 98.34 | 98.54 | 5 |
| MT-467 | 98.72 | 98.55 | 98.76 | 0 |
| MT-4854 | 98.59 | 98.4 | 98.63 | 3 |
| MT-4942 | 98.57 | 98.43 | 98.62 | 6 |
| MT-504 | 98.62 | 98.5 | 98.67 | 6 |
| MT-5195 | 98.54 | 98.38 | 98.6 | 2 |
| MT-5383 | 98.72 | 98.54 | 98.76 | 1 |
| MT-5488 | 98.76 | 98.62 | 98.79 | 0 |
| MT-5531 | 98.59 | 98.4 | 98.63 | 0 |
| MT-5543 | 98.52 | 98.34 | 98.57 | 2 |
| MT-567 | 98.70 | 98.54 | 98.74 | 2 |
| MT-578 | 98.79 | 98.68 | 98.84 | 0 |
| MT-5983 | 98.76 | 98.59 | 98.81 | 2 |
| MT-6084 | 98.51 | 98.37 | 98.56 | 3 |
| MT-6218 | 98.00 | 97.69 | 98.06 | 0 |
| MT-6226 | 98.65 | 98.45 | 98.7 | 0 |
| MT-6429 | 98.63 | 98.46 | 98.67 | 2 |

^a^ Only three lanes available for this sample.

**Table S2**. Comparison of consensus single-nucleotide polymorphisms (cSNPs) and heterogeneous alleles (hSNPs) in all samples aligned to H37Rv versus MT-0080_PB, after initial filtering with Allelic Fraction for cSNPs ≥ **0·95** and **0·05** < hSNP < **0·95**

|  | **cSNPs in all 62 samples** | | | | **hSNPs in all 62 samples** | | | |
| --- | --- | --- | --- | --- | --- | --- | --- | --- |
|  | **H37Rv reference, n=51430** | | **MT-0080 reference, n=365** | | **H37Rv reference, n=4897** | | **MT-0080 reference, n=125** | |
|  | **Median (IQR)** | **Range** | **Median (IQR)** | **Range** | **Median (IQR)** | **Range** | **Median (IQR)** | **Range** |
| Phred | 24922·77 (12084·77, 29426·77) | **596·85,** 55420·77 | 28115·77 (25306·77, 31561·77) | 792·77, 47999·77 | 3820·76 (1065·77, 8864·77) | 50·77, 37538·77 | 136·77 (84·77, 1872·77) | 50·77, 26808·77 |
| RMS-MQ | 60 (60, 60) | **39·13**, 69·42 | 60 (60, 60) | 39·75, 60 | 58·91 (56·62, 59·95) | 33·81, 65·42 | 60 (60, 60) | 46·61, 60 |
| DP | 670 (322, 783) | 20, 1468 | 758 (683, 844) | 23, 1235 | 801 (421, 1056) | 20, 1500* | 256 (78, 687) | 32, 990 |
| ReadPosRankSum | -0·064 (-1·093, 1·043) | **-5·269**, 3·735 | 010 (-0·783, 0·913) | -2·84, 2·71 | 1·383 (-2·691, 5·418) | -7·992, 15·309 | 0·892  (-0·501, 2·502) | -6·366, 4·101 |
| FS | 0 (0, 0) | 0, **45·053** | 0 (0, 0) | 0, 2·28 | 9·28 (1·979, 26·173) | 0, 59·985 | 1·723 (0·685, 4·69) | 0, 58·993 |

Legend. Initial filtering thresholds used were: Phred score < **50**, Root Mean Square Mapping Quality [RMS-MQ] ≤ **30**, depth [DP] < **20**, Read Position Rank Sum [ReadPosRankSum] < **-8**, Fisher Strand Bias [FS] ≥ **60**. As a consequence of the depth of coverage, where allelic fraction was **0·05** < alternative allele [ALT] < **0·95**, all hSNPs had at least 2 REF and ALT alleles by default. This includes all hSNPs and cSNPs identified across all samples, except variants in PE_PGRS and PPE genes, as well as those in mobile elements; some of these variants will be in positions that are excluded from the core alignment, as they failed quality control or are missing in at least one sample in the dataset. Read Position Rank Sum can only be calculated when both reference and alternative alleles are present at a position, therefore the number of cSNPs included in the summary statistics for this variable are 14720 for the H37Rv alignment and 153 for the alignment to MT-0080. *As samples were downsampled to this threshold, this is truncated at 1500.

**Table S3.** Assembly metrics for Single Molecule Real-Time sequencing of MT-0080 (‘MT-0080_PB’), aligned to NC_000962.3 (H37Rv).

|  | **MT-0080_PB** |
| --- | --- |
| Number of contigs | 1 |
| Largest contig, length in bp | 4,426,525 |
| GC content, % | 65·61 |
| Genome fraction covered, % | 99·28 |
| Largest alignment, bp | 592,812 |
| Total aligned length, bp | 4,387,983 |
| NG50 | 4,426,525 |
| NA50 | 256,105 |
| Reads mapped, % | 97·48 |
| Average depth of coverage, using filtered reads | 235 |
| Coverage ≥ 10x, % | 100 |
| Number of relocations | 50 |
| Number of inversions | 1 |
| Number of missing bases (‘N’) | 0 |
| Number of CDS | 4,321 |
| Number of RNA | 47 |
| Average nucleotide identity to H37Rv | 99·92% |

Legend. Quast (v.5.0.2 ^19^) was used to tabulate the above statistics, with the exception of the number of CDS and RNA, where annotation was done using RASTtk^20^ (v.2.0).

**Table S4**. Comparison of consensus single-nucleotide polymorphisms (cSNPs) and heterogeneous alleles (hSNPs) in all samples aligned to H37Rv versus MT-0080_PB, after initial filtering with Allelic Fraction for cSNPs ≥ **0·99** and **0·01** < hSNP < **0·99**

|  | **cSNPs in all 62 samples** | | | | **hSNPs in all 62 samples** | | | |
| --- | --- | --- | --- | --- | --- | --- | --- | --- |
|  | **H37Rv reference, n=49965** | | **MT-0080 reference, n=361** | | **H37Rv reference, n=5819** | | **MT-0080 reference, n=129** | |
|  | **Median (IQR)** | **Range** | **Median (IQR)** | **Range** | **Median (IQR)** | **Range** | **Median (IQR)** | **Range** |
| Phred | 25180·77 (15127·77, 29568·77) | 610·77, 55420·77 | 28133·77 (25313·77, 31561·77) | 792·77, 47999·77 | 4457·77 (1338·53, 8490·77) | 50·77, 37538·77 | 157·77 (84·77, 1986·77) | 50·77, 43450·77 |
| RMS-MQ | 60 (60, 60) | 39·13, 69·31 | 60 (60, 60) | 39·75, 60 | 59·41 (56·64, 60) | 33·81, 69·25 | 60 (60, 60) | 46·61, 60 |
| DP | 676 (415, 787) | 20, 1468 | 759 (683, 844) | 23, 1235 | 664 (233, 1017) | 20, 1500* | 271 (79, 691) | 32, 1191 |
| ReadPosRankSum | 0·032 (-0·975, 1·09) | -2·898, 2·82 | 0·118 (-0·738, 0·953) | -1·868, 2·71 | 0·628 (-2·273, 4·538) | -7·992, 15·309 | 0·773 (-0·624, 2·475) | -6·366, 4·101 |
| FS | 0 (0, 0) | 0, 9·514 | 0 (0, 0) | 0, 0 | 5·933 (1·018, 22·791) | 0, 59·985 | 1·623 (0, 4·676) | 0, 58·993 |

Legend. Initial filtering thresholds used were: Phred score < **50**, Root Mean Square Mapping Quality [RMS-MQ] ≤ **30**, depth [DP] < **20**, Read Position Rank Sum [ReadPosRankSum] < **-8**, Fisher Strand Bias [FS] ≥ **60**. Where **0·01** < alternative allele [ALT] < **0·99**, a minimum of 2 REF/ALT alleles were required for all hSNPs to reduce risk of including sequencing error; those that failed to meet these criteria were excluded. This includes all hSNPs and cSNPs identified across all samples, except variants in PE_PGRS and PPE genes, as well as those in mobile elements; some of these variants will be in positions that are excluded from the core alignment, as they failed quality control or are missing in at least one sample in the dataset. Read Position Rank Sum can only be calculated when both reference and alternative alleles are present at a position, therefore the number of cSNPs included in the summary statistics for this variable are 13255 for the H37Rv alignment and 148 for the alignment to MT-0080 in the cSNPs ≥ **0·99** and **0·01** < ALT < **0·99** analysis. *As samples were downsampled to this threshold, this is truncated at 1500. P values were calculated using on the Wilcoxon-Mann-Whitney test.

**Figures.**

**Figure S1.** Pileup of reads showing hSNPs suspected to be due to alignment error as listed in **Supplemental Dataset 1**, with MT-4942 used as an example and zoomed on position 2,255,171 to 2,280,170 in H37Rv (National Center for Biotechnology Information RefSeq Database Accession NC_000962.3).

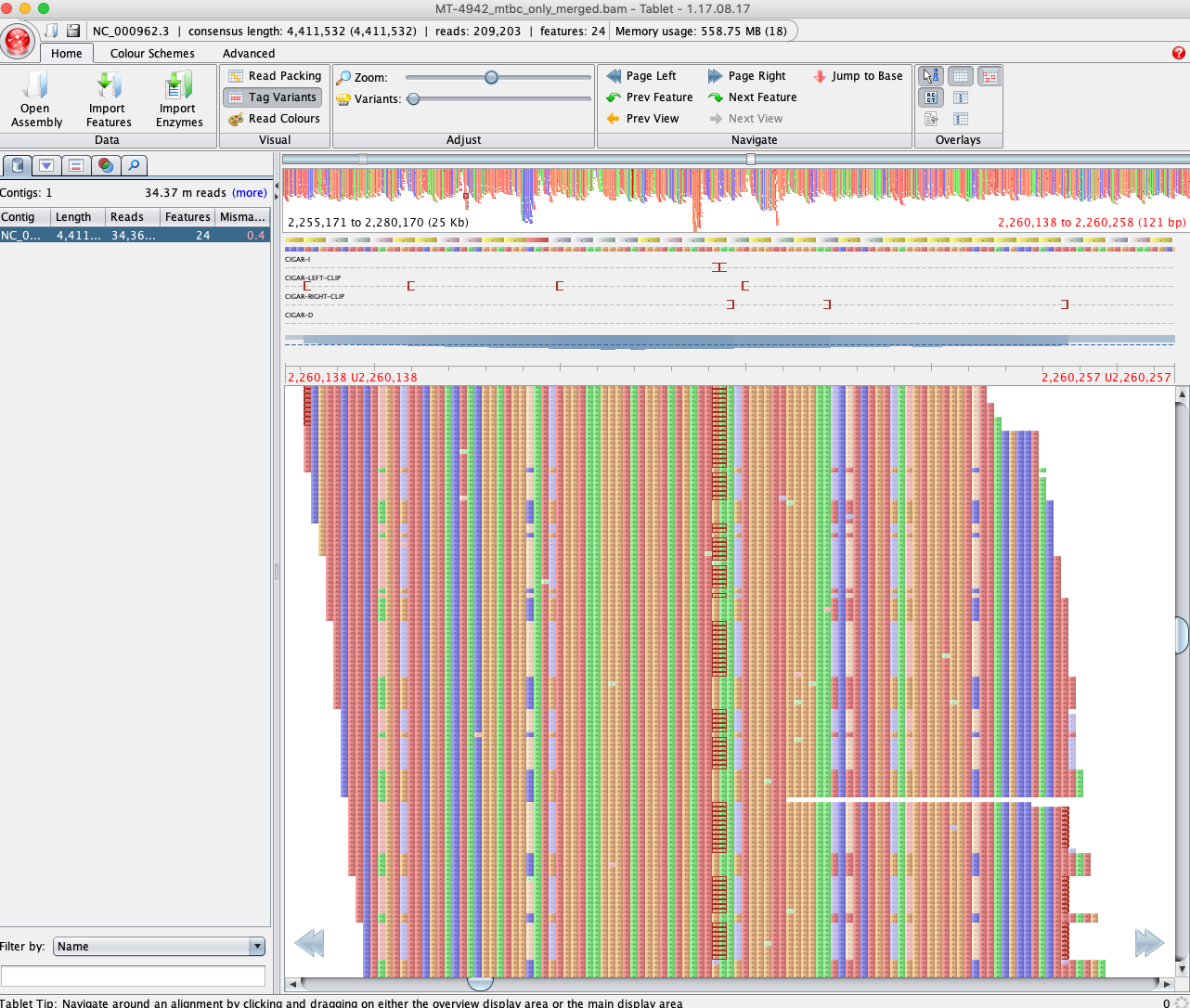

Legend. Binary Alignment Map (BAM) file were loaded into Tablet (v.1.17.08.17, ^25^) to visualize the pileup compared to H37Rv.

**Supplementary Datasets**

**Supplemental Dataset 1.** hSNPs shared across all 62 samples compared to H37Rv.

Quality metrics are given for each hSNP present in all 62 samples, as well as average depth of coverage for each sample.

**Supplemental Dataset 2.** Core alignments to MT-0080_PB using different filtering criteria; positions where all samples share the same allele as the reference are not indicated and positions with missing/low-quality data in any sample are excluded. Core alignments are provided for each main filtering protocol compared; descriptions are provided on an extra datasheet. Samples are organized within their respective clusters (as identified by hierarchical Bayesian Analysis of Population Structure), and within these, sub-groups are coloured based on previous cSNP/epidemiological analysis. ^2^ Original sub-lineage names are indicated on the top row. Approximate date of diagnosis is indicated, and summary totals of cSNPs and hSNPs at each site are tabulated.

References.

12.	Page AJ, Taylor, B., Delaney, A.J., Soares, J., Seemann, T., Keane, J.A., Harris, S.R. SNP-sites: rapid efficient extraction of SNPs from multi- FASTA alignments. *Microb Genom* 2016.
